# Highly efficient transgenesis with *miniMos* in *Caenorhabditis briggsae*

**DOI:** 10.1101/704569

**Authors:** Qiutao Ding, Xiaoliang Ren, Runsheng Li, Luyan Chan, Vincy WS Ho, Yu Bi, Dongying Xie, Zhongying Zhao

## Abstract

*C. briggsae* as a companion species for *C. elegans* has played an increasingly important role in study of evolution of development, gene regulation and genome. Aided by the isolation of its sister spices, it has recently been established as a model for speciation study. To take full advantage of the species for comparative study, an effective transgenesis method especially those with single copy insertion is important for functional comparison. Here we modified a transposon-based transgenesis methodology that had been originally developed in *C. elegans* but worked marginally in *C. briggsae*. By incorporation of a heat shock step, the transgenesis efficiency in *C. briggsae* with single copy insertion is comparable to that in *C. elegans*. We used the method to generate 54 independent insertions mostly consisting of a mCherry tag over the *C. briggsae* genome. We demonstrated the use of the tags in identifying interacting loci responsible for hybrid male sterility between *C. briggsae* and *C. nigoni* when combined with the GFP tags we generated previously. Finally, we demonstrated that *C. briggsae* has developed native immunity against the *C. elegans* toxin, PEEL-1, but not SUP-35, making the latter a potential negative selection marker against extrachromosomal array.

**Summary:** Nematode *C. briggsae* has been used for comparative study against *C. elegans* over decades. Importantly, a sister species has recently been identified, with which *C. briggsae* is able to mate and produce viable hybrid progeny. This opens the possibility of using nematode species as a model for speciation study for the first time. To take full advantage of *C. briggsae* for comparative study, an effective transgenesis method to generate single copy insertion is important especially for functional comparison. An attempt was made previously to generate single copy insertion with transposon-based transgenesis methodology, which had been originally developed in *C. elegans* but with limited success in *C. briggsae*. Here we modified the transposon-based methodology by incorporation of a heat shock step, which allows us to achieve a much higher transgenesis efficiency in *C. briggsae* with single copy insertion. We used the method to generate 54 independent insertions mostly consisting of a mCherry tag over the *C. briggsae* genome. We demonstrated the use of the tags in identifying interacting loci responsible for hybrid male sterility between *C. briggsae* and *C. nigoni* when combined with the GFP tags we generated previously. Finally, we demonstrated that *C. briggsae* has developed native immunity against the *C. elegans* toxin, PEEL-1, but not SUP-35, making the latter a potential negative selection marker against extrachromosomal array. Taken together, the modified transgenesis methodology and the transgenic strains generated in this study are expected to further facilitate *C. briggsae* as a model for comparative study or speciation study.

## Introduction

As a comparative model of *Caenorhabditis elegans*, *C. briggsae* shares a similar morphology, carries a genome of comparable size (Ren et al., 2018; Ross et al., 2011; Stein et al., 2003; Yin et al., 2018) and adopts similar developmental pattern to that of *C. elegans* (Zhao et al., 2008). As a model organism, *C. elegans* genome has been subject to intensive manipulations with various tools, including random mutagenesis with ultraviolet light coupled with trimethylpsoralen (UV/TMP) (Thompson et al., 2013), low copy insertion of transgenes with biolistic bombardment (Hochbaum, Ferguson, & Fisher, 2010; Praitis, Casey, Collar, & Austin, 2001; Radman, Greiss, & Chin, 2013), targeted single copy insertion using transcription activator-like effector nucleases (TALEN) system (Lo et al., 2013; Wood et al., 2011), or using CRISPR/Cas9 system (Chiu, Schwartz, Antoshechkin, & Sternberg, 2013; Cho, Lee, Carroll, Kim, & Lee, 2013; Dickinson & Goldstein, 2016; Dickinson, Pani, Heppert, Higgins, & Goldstein, 2015; Waaijers et al., 2013), *Mos1*-mediated single-copy insertion (MosSCI) (Frøkjær-Jensen et al., 2008) as well as random single-copy insertion with *miniMos* (Frøkjær-Jensen et al., 2014). Many of these tools have been adopted in *C. briggsae* for genome editing with a comparable or a much-reduced successful rate. For example, biolistic bombardment was successfully adopted in *C. briggsae* transgenesis with comparable efficiency, whereas single-copy insertion with *miniMos* demonstrated a much lower efficiency in *C. briggsae* than in *C. elegans* (Bi et al., 2015; Frøkjær-Jensen et al., 2014; Zhao et al., 2010). An efficient transgenesis method with single copy insertion is essential for comparative functional study of biology between *C. elegans* and *C. briggsae*. Recent work in *Caenorhabditis* species has demonstrated that heat shock treatment or compromised function in heat shock pathway significantly increased the frequency of transposition (Ryan, Brownlie, & Whyard, 2016), raising the possibility of improving *miniMos*-based transgenesis in *C. briggsae* by incorporation of a heat shock step or inhibition of activities of heat shock proteins.

Isolation of *C. briggsae* sister species, *C. nigoni*, with which it can mate and produce viable progeny, paves the way of speciation study using nematode as a model for the first time (Kozlowska, Ahmad, Jahesh, & Cutter, 2012; Woodruff, Eke, Baird, Félix, & Haag, 2010). To isolate post-zygotic hybrid incompatible (HI) loci between *C. briggsae* and *C. nigoni*, a dominant and visible marker is required for targeted introgression, in which the genome of one species is labeled with the marker and repeatedly backcrossed into the other to isolate the HI loci. The marker greatly facilitates tracing of its linked genomic fragment during backcrossing and its associated HI phenotype. Given numerous such markers are required over a genome, generation of such markers using targeted single-copy insertion becomes an enormous burden. This is because that the efficiency for targeted single-copy insertion is relatively low. As an alternative, over 100 GFP markers were inserted into the *C. briggsae* genome with biolistic bombardment, which allowed genome-wide mapping of HI loci between the two species (Bi et al., 2015; Ren et al., 2018; Yan, Bi, Yin, & Zhao, 2012).

Biolistic bombardment using *cbr-unc-119* (Bi et al., 2015; Zhao et al., 2010) generates low copy number of transgene into genome but requires tedious animal preparations. It also needs to wait for several weeks before screening for transformed animal. In addition, presumably due to dramatic mechanic shearing, the transgenic animals often suffer from chromosomal rearrangements (Bi et al., 2015; Ren et al., 2018; Tyson et al., 2018). Importantly, the insertion site cannot be precisely determined due to unknown copy number of the transgene and its arrangement within host genome. As such, the transgene insertion sites were estimated by genotyping introgression boundary with PCR using species-specific primers (Bi et al., 2015; Yan et al., 2012). Unfortunately, the divergence between the *C. briggsae* and *C. nigoni* genomes is too far to allow efficient recombination during backcrossing, resulting in a poor mapping resolution of the insertion site (Bi et al., 2019, 2015; Yan et al., 2012).

We have recently demonstrated that inter-chromosomal interactions are involved in hybrid male sterility between *C. briggsae* and *C. nigoni* (Bi et al., 2019; Li et al., 2016). For example, a *C. briggsae* X chromosome fragment in an otherwise *C. nigoni* background as an introgression leads to male sterility. Hybrid F1 male sterility can be rescued by the presence of the introgression and another *C. briggsae* genomic fragment on its Chromosome II. We speculate that such interacting loci might be common in producing HI. Availability of the dual-color labeling system makes it possible for systematic mapping such interacting loci. This is because it facilitates tracking of co-segregation of the two loci. It is worth noting that most existing markers in the *C. briggsae* genome were derived from GFP, preventing effective screening for such loci. Therefore, *C. briggsae* animals bearing a visible marker other than the GFP are desired.

Transposon-mediated single copy insertion at random genomic position with high efficiency has been successfully developed in *C. elegans* (Frøkjær-Jensen et al., 2014). A fly transposon *Mos1* was truncated with minimal transposon sequences, termed as *miniMos* cargo vector, in order to boost its capacity of carrying foreign DNA (Frøkjær-Jensen et al., 2014). Single copy of transgene can be inserted into host genome at high frequency via co-injecting a transposase-expressing vector and *miniMos* cargo vector into *C. elegans* gonad. The method was also adopted in *C. briggsae* but with a much lower efficiency with unknown reasons (Frøkjær-Jensen et al., 2014). Attempts have been made to improve insertion frequency in *C. briggsae*. For example, the promoter (*cel*-P*eft-3*) driving transposase expression was substituted with *C. briggsae* promoter *cbr*-P*eft-3* or *cbr*-P*pie-1* in order to boost transposase expression. However, the insertion frequency did not improve as expected. A highly efficient method for inserting single-copy transgene has yet to be established in *C. briggsae*. To facilitate screen for single-copy insertion, a negative selection marker *cel*-P*hsp-16.41*∷*peel-1* was co-injected into animals to kill extrachromosomal array-bearing worms during screening (Frøkjær-Jensen, Davis, Ailion, & Jorgensen, 2012). However, the killing efficiency of the negative selection marker *peel-1* has not been investigated in *C. briggsae*.

This study established a highly efficient methodology for inserting single-copy transgene into the *C. briggsae* genome based on *miniMos*. Negative selection marker against transgene consisting of extrachromosomal array was also explored with limited success.

## Materials and Methods

### Nematode strains

*C. elegans*, *C. briggsae* and *C. nigoni* wild isolates used in this study were N2, AF16 and JU1421, respectively. The *C. briggsae dpy-5(zzy0580)* knockout strain ZZY0580 was generated using CRISPR/Cas9 with AF16 followed by backcrossing to AF16 for three generations. Introgression strains used were ZZY10330 (*zzyIR10330* (*X* [*cbr-myo-2p∷gfp*, *cbr-unc-119(*+)]), AF16>JU1421) (Bi et al., 2015), ZZY10377 (*zzyIR10377* (*X* [*cbr-myo-2p∷mCherry*, neoR]), AF16>JU1421) from *C. briggsae* transgenic strain ZZY0777 (Table S2). Introgression strains ZZY10353 (*zzyIR10353* (*II* [*cbr-myo-2p∷gfp*, *cbr-unc-119(*+)]), AF16>JU1421) (Bi et al., 2015) and ZZY10382 (*zzyIR10382* (*II* [*cbr-myo-2p∷mCherry*, neoR]), AF16>JU1421) were generated from *C. briggsae* transgenic strain ZZY0782 (Table S2). Details for all the transgenic strains generated in this study were included in Table S2. All the *C. elegans* and *C. briggsae* lines were maintained on 1.5% nematode growth medium (NGM) seeded with *E. coli* OP50 at room temperature and in a 25 °C incubator, respectively, unless specified otherwise.

### Introgression

Introgression was performed as described (Bi et al., 2015). Specifically, *C. briggsae* transgenic strains, ZZY0777 and ZZY0782 each expressing a single-copy of P*myo-2*∷mCherry located on the X and chromosome II, respectively, were backcrossed to *C. nigoni* (JU1421) for 15 generations to give rise to ZZY10377 and ZZY10382. Introgression boundaries were mapped as described (Yan et al., 2012).

### Molecular Biology and transgenesis

pZZ203 was derived from pZZ0031 (Yan et al., 2012) by removing *cbr*-P*myo-2∷gfp∷his-72* UTR by cutting using KpnI and ApaI (NEB). The digested pZZ0031 backbone carrying *cbr-unc-119*(+) was gel purified, blunt end repaired, and self-ligated to give rise to pZZ203. pZZ160 and pZZ161 was derived from pCFJ910[NeoR] and pCFJ909[*cbr-unc-119*(+)], respectively, by inserting both *cbr*-P*myo-2∷gfp∷his-72* UTR and *cbr-dpy-5*(+) into minimal *Mos1* transposon. pZZ184 was derived from the pZZ160 by replacing *gfp∷his-72 UTR* and *cbr-dpy-5(+)* with *mCherry∷unc-54* UTR from pGH8 (Frøkjær-Jensen et al., 2008). The plasmids pZZ185 and pZZ196 for negative selection were derived from pCFJ601[P*eft-3*∷Mos1 transposase∷*tbb-2* UTR] by replacing the Mos1 with *sup-35* (Ben-David, Burga, & Kruglyak, 2017) and *peel-1* (Seidel et al., 2011; Seidel, Rockman, & Kruglyak, 2008), respectively. The plasmid kit containing pCFJ601, pCFJ909, pCFJ910, pGH8, and pMA122 was acquired from Addgene (cat# 1000000031). Vector pZZ113 containing sgRNA expression cassette against *cbr-dpy-5* was derived from P*U6*∷*unc-119*_sgRNA (Addgene plasmid # 46169) as described (Friedland et al., 2013). The *cbr-dpy-5* gRNA target sequence is GGAGCCCCAGGAGAGCCAGG. Primers used for construct building were listed in Table S1. An overview of plasmid compositions was shown in Fig. S1.

Plasmids for microinjection were extracted using the PureLink HQ Mini Plasmid Purification Kit or HiPure Plasmid Midiprep kit (Invitrogen). Genomic DNAs were extracted from mix-staged animals with PureLink Genomic DNA Mini Kit (Invitrogen). Injection mixture for transformation consisted of *miniMos* cargo vector at 20 ng/μl, *Mos1* transposase-expressing plasmid at 50 ng/μl, red or green fluorescent co-injection markers each at 10 ng/μl. pZZ203 was added to 80 ng/μl with a total DNA concentration of 170 ng/μl. Plasmids pMA122, pZZ185, and pZZ196 were built to test their use as a negative selection marker. Neomycin stock solution of 12.5 mg/ml was made with G418 powder (ThermoFisher) in water. For detailed transformation steps, see Fig. 1.

**Fig.1.**
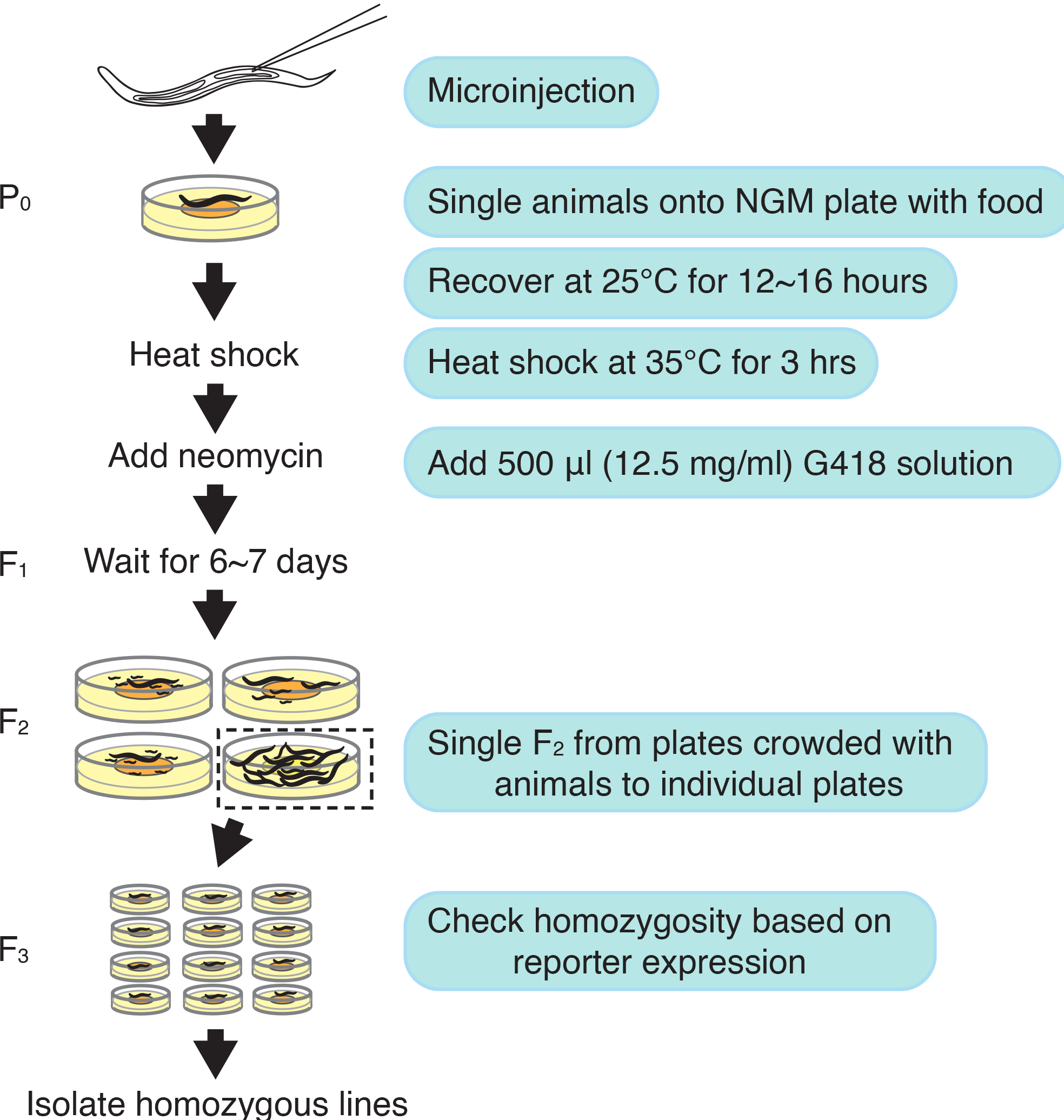
Schematics of optimized protocol for *miniMos*-based transgene insertion in *C. briggsae*. Injected animal was individually placed on a 55 mm crossing plate seeded with OP50 (~6.7 ml NGM per plate) and allowed to recover at 25 °C overnight (roughly 12 to 16 hrs). For heat shock treatment, plates with injected animals were sealed with parafilm and incubated on the water bath pre-heated to 35 °C for 3 hrs. Remove parafilm and any water droplet on the inner lid. Incubate plates at room temperature (approximately 22 °C) for half an hour. Add 0.5 ml 12.5 mg/ml neomycin (G418) solution pre-warmed up at room temperature onto each plate. Gently rotate plates to allow neomycin solution to fully cover plate surface. Air-dry the plates and incubate the dried plates in a 25 °C incubator for 6-7 days. Due to the presence of selection marker NeoR in the injected *miniMos* plasmids (pZZ160 and pZZ184), single 10 to 12 survived L4 or young adult animals from the plates crowded with animals that do not express co-injection marker onto individual plates to be incubated at 25 °C. After 2-3 days, harvest animals with homozygous transgene based on the reporter expression and lack of expression of visible co-injection marker for genotyping.

### Quantification of P_0_ insertion frequency

Three days after microinjection, plates without F_1_ transgenic progeny were discarded and excluded from the number of injected P_0_ animals. The F_2_ animals were screened for *miniMos* insertion under stereo-microscope based on reporter expression or phenotypic rescue, and the absence of co-injection markers. Animals confirmed with transgene insertion from the same P_0_ plate were treated as a single independent insertion. The P_0_ insertion frequency was measured through dividing the number of independent insertions by the total number of injected P_0_ animals. Mapping of insertion site was performed as described (Frøkjær-Jensen et al., 2014). The *C. briggsae* cb4 genome (Ross et al., 2011) was used to report the chromosomal coordinates.

### Selection marker for transformation

For selection with Neomycin (NeoR), injected animals were allowed to recover overnight (12 to 16 hours) followed by heat shock treatment on a 35 °C water bath for three hours. The heat shock treated plates were incubated at room temperature for half hour to recover. Then 500 μl of 12.5 mg/ml G418 solution was added to each plate. Plates were incubated at 25 °C for 6 to 7 days. Twelve F_2_ animals were singled onto a new NGM plate from the P_0_ plates crowded with animals. Homozygosity was checked based on reporter expression in F_3_.

For selection with *cbr-dpy-5*, all rescued *dpy-5*(+) F_1_ animals were singled onto individual plates. Twelve F_2_ animals were singled onto new plate from each F_1_ plate in which over 50% animals were *dpy-5*(+) ones. F_2_ plates with 100% *dpy-5*(+) animals that did not express visible co-injection markers were kept for genotyping.

### Development of negative selection marker for extrachromosomal array

A fusion PCR product was made between a cassette consisting of *peel-1*∷*tbb-2* UTR or *gfp*∷*tbb-2* UTR and the *C. briggsae* syntenic region of *hsp-16.41* promoter with the primers listed in Table S1. Additional details of *C. briggsae* syntenic region of *hsp-16.41* promoter sequence screening are available in Fig. S2. Injection mixture was made with pMA122, pZZ185, pZZ196 or the fusion PCR product at 70 ng/μl, fluorescent co-injection marker vector pZZ161 at 20 ng/μl, pGH8, pCFJ90, pCFJ104 at 10 ng/μl each, and pZZ203 at 50 ng/μl.

To examine the lethality in *C. briggsae* after heat shock, an array-bearing line was generated and embryos harvested for synchronization. The synchronized L1 animals were divided into two groups, one was subjected to heat shock treatment at 35 °C for 3 hours before transferring into 25 °C incubator, and the other was incubated at 25 °C without heat shock treatment as a control. *C. elegans* heat shock treatment was performed at 34 °C for 3 hours. The ratio of surviving adults carrying an array out of all progeny were calculated in both control and heat-shock treated animals.

## Results

### Heat shock treatment significantly increased the efficiency of transgene insertion

To improve the transgenesis efficiency using *miniMos* in *C. briggsae*, we started the transgenesis by repeating the steps basically as described previously (Frøkjær-Jensen et al., 2014), i.e., injection of a vector carrying *cbr*-P*myo-2*∷*gfp* along with a transposase-expressing vector into *cbr-unc-119* deletion mutant strain RW20000 (Zhao et al., 2010). One of major complications associated with the *cbr-unc-119* as an injection selection marker was the low transformation efficiency even for the formation of extrachromosomal array. This was the case using the rescuing fragment from either *C. elegans* or *C. briggsae* (data not shown). We speculated that the low insertion rate associated with the selection marker could be related to the low transformation efficiency of single copy insertion.

To improve transformation efficiency, we generated another injection selection marker *cbr-dpy-5* using CRISPR/Cas9. This was based on the observation that *dpy-5* had been used as a very efficient selection marker in *C. elegans* microinjection (Hunt-Newbury et al., 2007; Zhao, Sheps, Ling, Fang, & Baillie, 2004). As expected, *cbr-dpy-5* worked as an effective selection marker for microinjection. We successfully obtained three independent transgenic lines with single copy insertion using the marker after injecting 35 animals. However, due to lack of negative selection maker against extrachromosomal array, it is tedious to screen for the rescued animals out of all progeny.

We then performed *miniMos-based* single-copy insertion using neomycin as a selection marker as described (Frøkjær-Jensen et al., 2014; Giordano-Santini et al., 2010). The method provided a key advantage in simplifying selection of the transformed animals because all un-rescued animals were killed, but the overall efficiency of obtaining single-copy insertion was not improved compared with previous studies (Fig. 2).

**Fig. 2.**
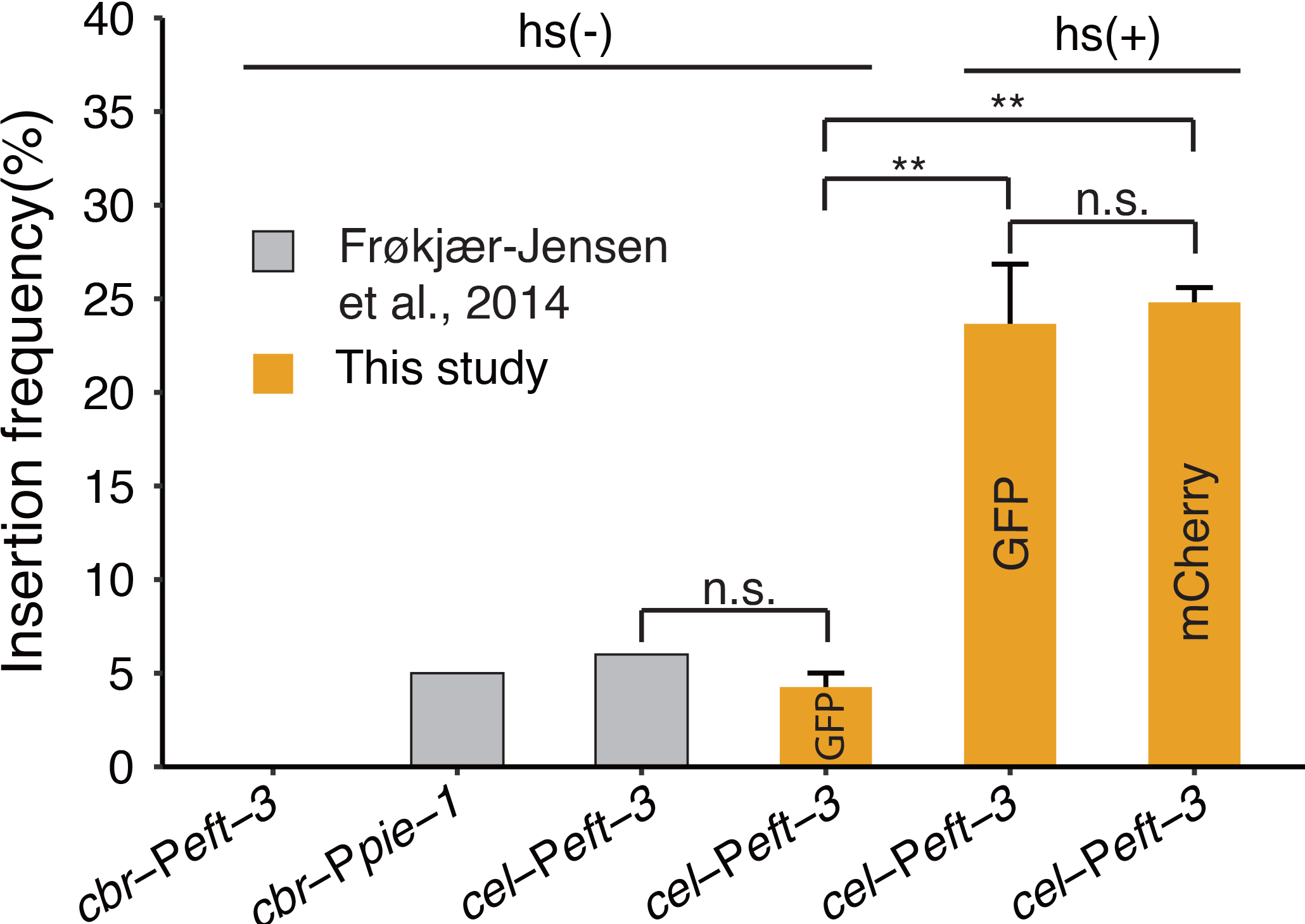
Comparison of insertion frequencies in *C. briggsae* between this (brown) and previous study (grey) (Frøkjær-Jensen et al., 2014). Y-axis denotes insertion frequency in P_0_ animals. X-axis indicates various promoters used in the previous and this study that drive Mos1 transposase expression. Each fluorescent marker in this study was tested for at least three replicates. Animals with or without heat shock are indicated with hs(+) and hs(−), respectively. Student’s t-test was performed for comparison between animals or without heat shock. Error bars represent standard error.

Given that heat shock treatment significantly increased transposition frequency (Ryan et al., 2016), we reasoned that inclusion of a heat shock step might boost the successful rate of transgenesis in *C. briggsae* based on *miniMos*, which is a modified transposon. As expected, inclusion of a step of heat shock treatment, i.e., 35 °C for 3 hrs, we were able to significantly increase the frequency of single-copy insertion of P*myo-2*∷*gfp* or P*myo-2*∷mCherry (Figs. 1 & 2). Insertion frequency increased from lower than 5% without heat shock to approximately 25% with heat shock for both constructs, indicating that the heat shock treatment was essential for increasing transposition rate.

### A large collection of single copy insertions expressing Pmyo-2∷mCherry were generated in C. briggsae

To complement the genetic resources mainly consisting of GFP insertions over the *C. briggsae* we generated previously (Bi et al., 2015), we decided to generate a cohort of insertions randomly distributed over the *C. briggsae* genome with a focus on generation of insertion expressing a fluorescent protein other than GFP, i.e., P*myo-2*∷mCherry. This would be particularly useful for isolation of interacting loci responsible for hybrid male sterility as detailed below. To this end, we generated a total of 54 insertions consisting of 9 P*myo-2*∷*gfp* and 45 P*myo-2*∷mCherry that were able to be uniquely mapped to the *C. briggsae* genome (Fig. 3 and Table S2). The markers show roughly random distribution over the *C. briggsae* genome.

**Fig. 3.**
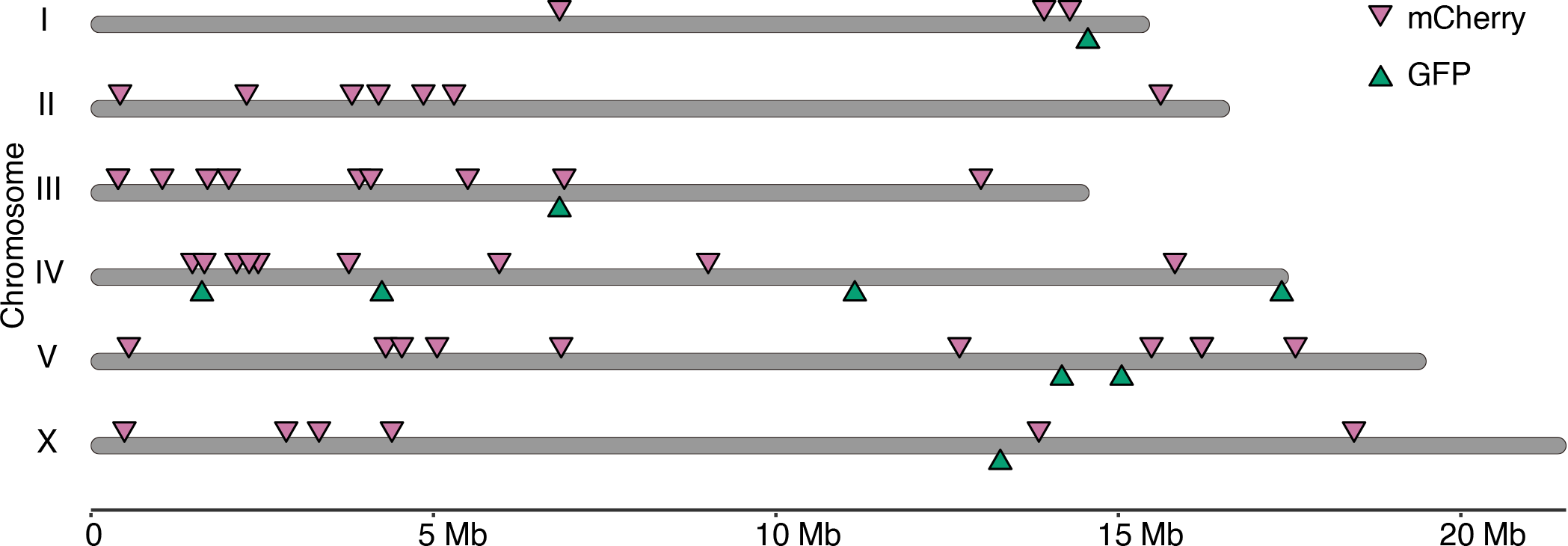
Genomic position of mapped insertions of visible marker P*myo-2*∷*gfp* or P*myo-2*∷mCherry over *C. briggsae* genome (cb4). Insertions landed onto contigs unassigned to a specific chromosome were not shown. Insertion*s* of individual GFP (green) or mCherry (purple) marker are indicated with inverted triangle along the genome in scale, detailed insertion information is listed in Table S2. Insertion site was determined using two rounds inverse PCR as described previously.

### Dual-coloured marking system facilitates screen for interacting loci responsible for HI loci

We demonstrated that genetic interaction between X Chromosome and autosome was essential for hybrid male sterility between *C. briggsae* and *C. nigoni* (Bi et al., 2019). Identification of such interacting loci is challenging without proper markers on interacting chromosomes. We showed that the interaction between independent GFP-labelled *C. briggsae* introgression fragments on two different chromosomes was essential for male fertility in *C. nigoni*, but it was difficult to pinpoint these interacting loci systematically. Availability of such dual-colour labelled strains would greatly facilitate the identification of such interactions. To help illustrate this point, we tried to recapitulate the interaction we identified earlier by using two *C. nigoni* strains ZZY10377 and ZZY10382 that carry a mCherry-labelled introgression fragment derived from the right arm of the *C. briggsae* X and chromosome II (Fig. 4), respectively, between which an interaction was found (Bi et al., 2019). We crossed two introgression strains, one labelled with GFP and the other with mCherry in reciprocal way. We then examined the fertility of the males that simultaneously carried both introgressions. We found that the males carrying the both loci were mostly fertile, whereas the males carrying a single introgression of GFP or mCherry inserted on the X chromosome were sterile, indicating that the dual-colour labelled introgressions did recapitulate the interaction. Presence of the dual-coloured markers paves the way for genome-wide identification of any other interacting loci responsible for the hybrid incompatibilities.

**Fig. 4.**
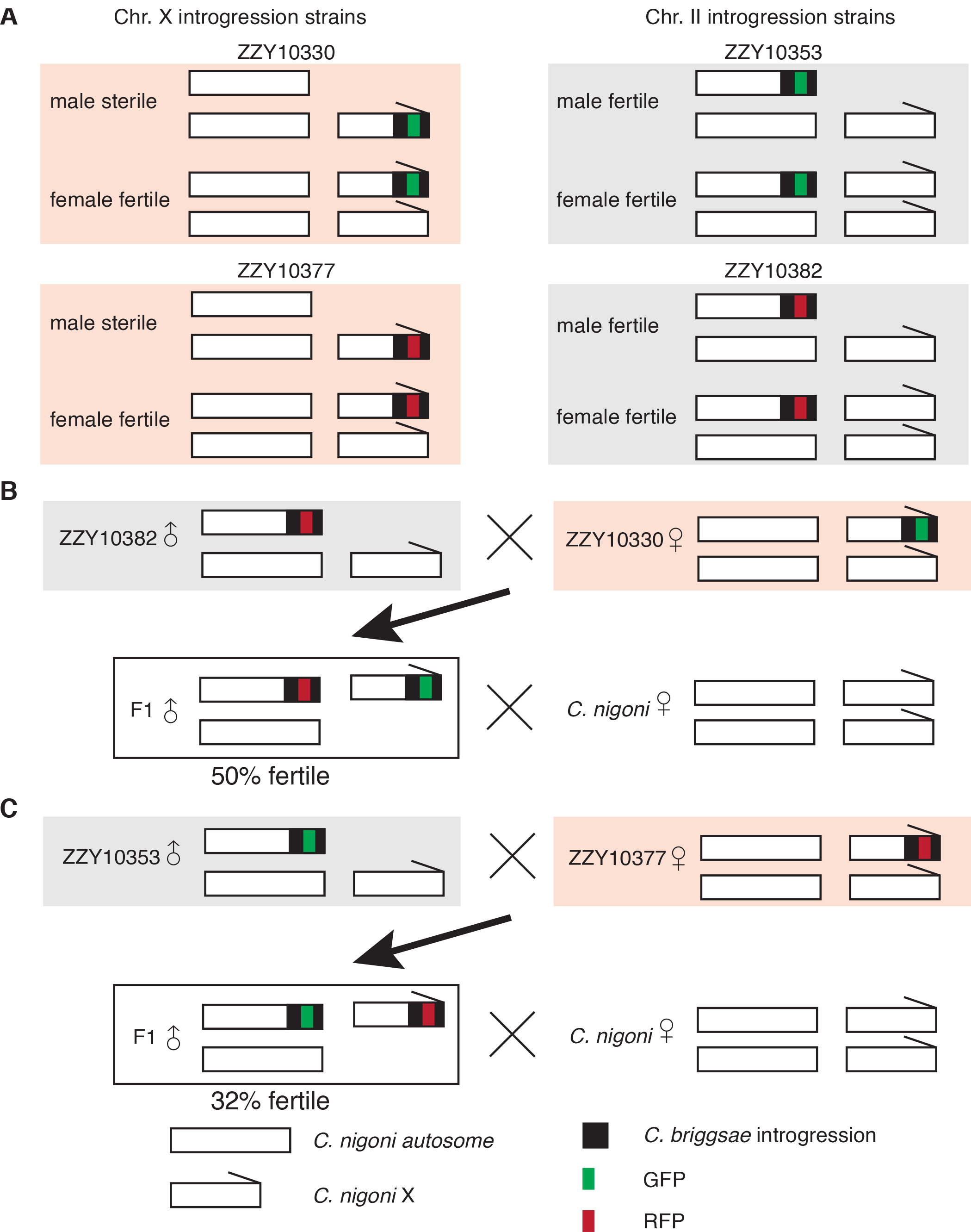
Use of dual-colour fluorescent markers in mapping interacting loci responsible for rescue of HI phenotype. A. Schematics of introgression strains expressing GFP marker (green) generated previously with biolistic bombardment or mCherry marker (red) generated with *miniMos* in this study. Strains ZZY10330 and ZZY10377 carry an X-linked introgression derived from *C. briggsae* that produces GFP or RFP expression, respectively, and HI phenotype in an otherwise *C. nigoni* background, i.e., hybrid male sterility. Strains ZZY10353 and ZZY10382 carry a Chromosome II-linked introgression derived from *C. briggsae* that produces GFP or RFP expression, respectively, and HI phenotype in an otherwise *C. nigoni* background, i.e., homozygous inviable (not shown). B. Schematics of crossing strategy in mapping interacting loci between X chromosome and Chromosome II. We previously demonstrated that presence of the introgression fragment from ZZY10353 rescued the male sterility of ZZY10330. However, this demands tedious genotyping of ZZY10353 to ensure its presence because it is impossible to distinguish the two introgressions both expressing GFP. Substitution of ZZY10353 with ZZY10382 expressing RFP greatly facilitated the process for screening for simultaneous presence of the two introgressions. C. Reciprocal crossing between strains with autosome- and X-linked introgressions expressing GFP and RFP, respectively, serves as the same purpose as in B.

### C. briggsae appeared to develop native immunity against PEEL-1

For transgene insertion using MosSCI or *miniMos* in *C. elegans*, a sperm-derived toxin gene, *peel-1*, is used as a negative selection marker to effectively kill extrachromosomal array-bearing worms (Frøkjær-Jensen et al., 2012, 2014; Seidel et al., 2011), greatly reducing the burden of screening for transgenic strains carrying an insertion out of all transgenic animals, including those carrying an extrachromosomal array.

To adopt the negative selection marker in *C. briggsae*, we first compared the killing effect of PEEL-1 in *C. elegans* and *C. briggsae* through its forced expression driven by a *C. elegans* heat shock promoter, P*hsp-16.41*. We generated transgenic lines carrying extrachromosomal array consisting of the PEEL-1-expressing vector in both *C. elegans* and *C. briggsae*. The synchronized L1 animals were subjected to heat shock at 34 °C and 35 °C for *C. elegans* and *C. briggsae*, respectively. The killing effect in *C. elegans* was comparable to that reported previously (Seidel et al., 2011), but no apparent killing was observed in *C. briggsae* after heat shock treatment (Fig. 5A).

**Fig. 5.**
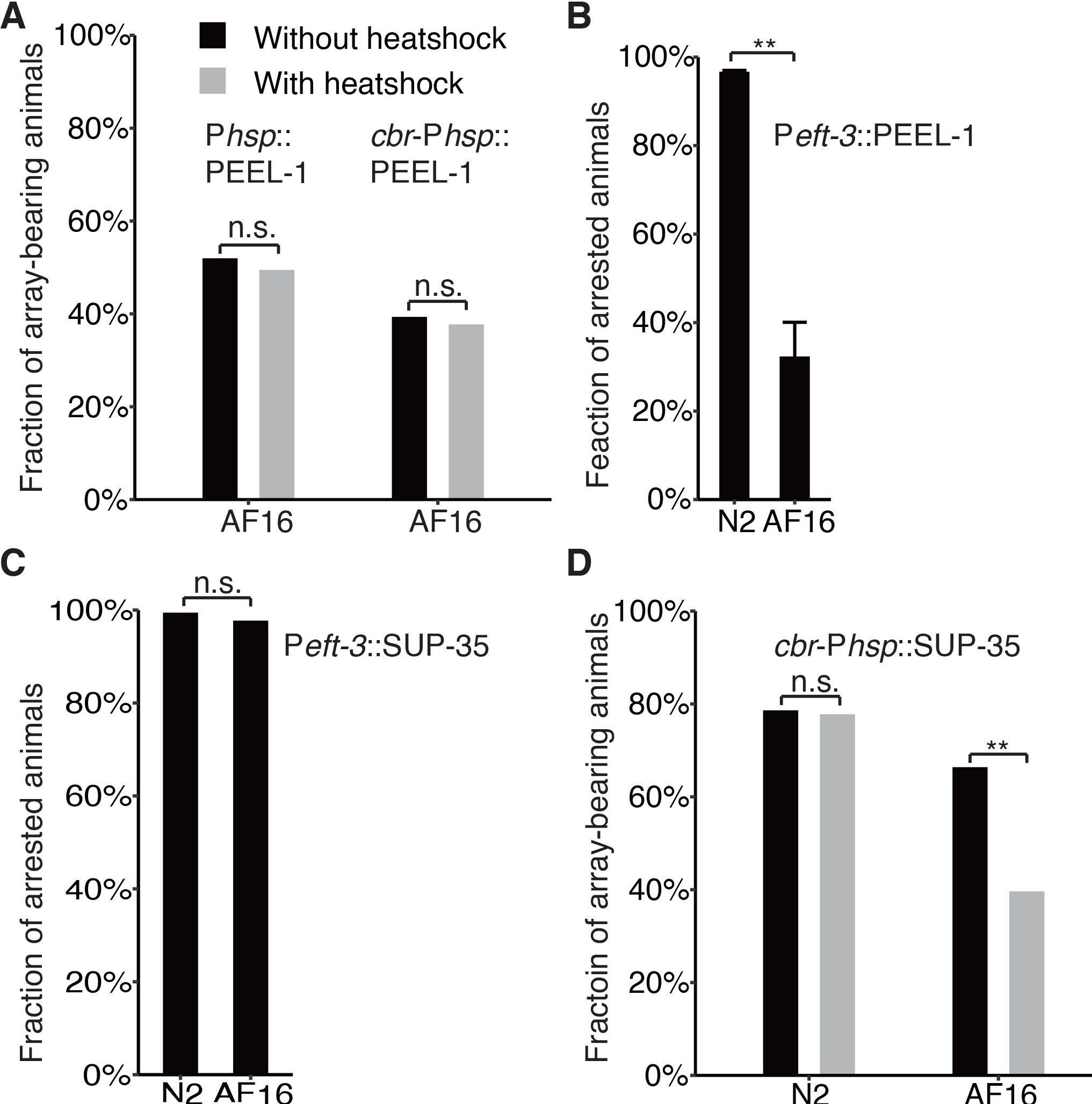
Comparison of killing efficiencies of PEEL-1 (A & B) and SUP-35 (C & D) in *C. elegans* (N2) and *C. briggsae* (AF16). (A) Fraction of surviving array-bearing animals with (grey) and without (black) heat shock in *C. briggsae* (AF16), which carries a *peel-1* driven by heat shock promoter as an extrachromosomal array as indicated. Note, no significant change in survival using *hsp-16.41* promoter of from *C. elegans* and CBG19187(*cbr-hsp-16.60*) promoter from *C. briggsae.* (B) Fraction of arrested F_1_ animals presumably carrying an injection marker derived from P_0_, which was injected with *peel-1* driven by a constitutively expressing promoter, P*eft-3*. Note the incomplete killing of PEEL-1 expressing animals in *C. briggsae*. (C) Fraction of arrested F_1_ animals presumably carrying a co-injection marker derived from P_0_, which was injected with *sup-35* driven by P*eft-3*. Note the complete killing of *sup-35* expressing animals in both *C. elegans* and *C. briggsae*. (D) Fraction of surviving array-bearing animals with (grey) and without (black) heat shock in *C. elegans* (N2) and *C. briggsae* (AF16), which carries a *sup-35* driven by a heat shock promoter from CBG19187(*cbr-hsp-16.60*) as an extrachromosomal array as indicated. Note, no significant change in survival in *C. elegans* and a significant drop in *C. briggsae* survival after heat shock.

To further investigate what caused the failure of PEEL-1 to kill *C. briggsae*, we replaced *C. elegans* heat shock promoter, P*hsp-16.41*, with its *C. briggsae* equivalent (Fig. S2) and generated the transgenic lines. Again, we did not observe significant increase in killing (Fig. 5A). We reasoned that the *C. briggsae* heat shock promoter might not be a functional equivalent of P*hsp-16.41*. To examine whether the *C. briggsae* syntenic heat shock promoter responds to heat shock treatment, we generated transgenic lines carrying extrachromosomal array consisting of a fusion between the *C. briggsae* heat shock promoter and GFP. We did see induced GFP expression in the transgenic animals after heat shock treatment in both *C. elegans* and *C. briggsae* (data not shown), suggesting that *C. briggsae* somehow develops immunity against PEEL-1. To further investigate *C. briggsae*’s immunity against PEEL-1, we generated transgenic *C. briggsae* animals expressing *peel-1* driven by an effective promoter, P*eft-3*, which was known to able to drive expression in *C. briggsae* (Frøkjær-Jensen et al., 2012, 2014). Again, we did not observe a significant killing effect (Fig. 5B). Taken together, it seems that *C. briggsae* develops native immunity against PEEL-1, preventing it from being used as a negative selection marker against extrachromosomal array.

### Limited success of using sup-35 as a negative selection marker for extrachromosomal array

In addition to *peel-1*, a maternal-effect toxin, SUP-35, that kills developing embryo, has recently been identified in *C. elegans* (Ben-David et al., 2017). To test the killing efficiency of SUP-35 in *C. briggsae*, we made a *sup-35*-expressing vector driven by the *C. elegans eft-3* promoter, P*eft-3* (Fig. S1). We injected the construct along with fluorescence co-injection markers into both *C. elegans* and *C. briggsae*. As expected, nearly all F_1_ embryos expressing the fluorescence markers arrested at late embryogenesis or as early larvae for both *C. elegans* and *C. briggsae* (Fig. 5C), indicating that SUP-35 is an effective toxin in *C. briggsae* and has a potential to be developed as a negative selection marker against extrachromosomal array in *C. briggsae*. To this end, we generated transgenic strains in *C. elegans* and *C. briggsae* that carry an extrachromosomal array, which consists of the *sup-35*-expressing vector driven again by *C. briggsae* equivalent of *hsp-16.41* promoter (Fig. S2). However, we did not observe any significant killing effect in *C. elegans* after heat shock treatment (Fig. 5D). Because *C. elegans* expresses both PEEL-1 and its antidote ZEEL-1 only transiently in embryo, but expresses SUP-35 and its antidote PHA-1 throughout its life cycle (Gerstein et al., 2010). It is possible that the postembryonic PHA-1 is sufficient to neutralize the toxicity of SUP-35 induced by heat shock treatment in *C. elegans*. It is also possible that the *C. briggsae* heat shock promoter may not respond to heat shock as effectively as its *C. elegans* equivalent in *C. elegans*. Consistent with this, the heat shock treatment showed a significant killing effect in *C. briggsae* carrying the same construct, i.e., from 66.4% to 39.6% though it is not efficient enough to serve as a negative selection marker.

## Discussion

Given the similar morphology, physiology and developmental program between *C. elegans* and *C. briggsae*, methods for *C. elegans* transgenesis are expected to be transferrable to *C. briggsae* with minimal modification. However, there is an exception to this, which is the case for *miniMos* mediated single-copy insertion (Frøkjær-Jensen et al., 2014).

### Potential cause of low efficiency of transgenesis in C. briggsae mediated by miniMos

It has been well established that transposon mobility can be significantly increased across species after exposure to biotic or abiotic stress (Bouvet, Jacobi, Plourde, & Bernier, 2008; Liu, Chu, Choudary, & Schmid, 1995; Walbot, 1999). Heat shock proteins serve as molecular chaperone to facilitate folding of other proteins into their appropriate conformations following heat exposure, thus buffering the subsequent phenotypic changes (Erlejman, Lagadari, Toneatto, Piwien-Pilipuk, & Galigniana, 2014). Perturbation of heat shock protein also increases transposition frequency in a similar way to that of heat treatment itself. It is possible that under optimal growth condition, *C. briggsae* has a more robust system to maintain genome integrity in its germline than *C. elegans*, which prevents its genome from environment-induced damage or editing. Improvement of transgenesis efficiency by heat shock treatment or inhibition of heat shock pathway could be at the cost of compromised genome integrity. One important way to preserve genome integrity in the germline is through Piwi-interacting RNA (piRNA), which constitutes one of the major regulatory molecules that curb the transposon activities (Ishizu, Siomi, & Siomi, 2012). Consistent with this, functional perturbation of Hsp90 attenuates the piRNA silencing mechanism, leading to transposon activation and the induction of morphological mutants in *Drosophila* (Specchia et al., 2010). Therefore, the transgenic strains generated using heat shock treatment should be subjected to rigorous backcrossing with wild isolate before any functional characterization.

### Differential responses to PEEL-1 and SUP-35 between C. elegans and C. briggsae

It is intriguing that *C. briggsae* responds differentially to the overexpressed paternal toxin PEEL-1 and maternal toxin SUP-35 from *C. elegans.* In *C. elegans*, PEEL-1 was believed suspected to function as a calcium pump on the cell membrane by generating a membrane pore, which leads to the release of intracellular calcium (Seidel et al., 2011). The maternal-effect toxin, SUP-35, kills developing embryos, but the molecular mechanism of its toxicity remains unclear (Ben-David et al., 2017). Our data showed that *C. briggsae* somehow develops native immunity against PEEL-1 but not SUP-35 (Fig. 5). However, forced expression of SUP-35 driven by the *C. briggsae* heat shock promoter did not mediate complete killing of *C. briggsae* after heat shock treatment (Fig. 5D). Two possible reasons account for the ineffective killing. First, time windows for SUP-35 killing seems to be narrow, i.e., at certain stage of embryogenesis. This is why a constitutive promoter, i.e., *eft-3* promoter is able to drive SUP-35 expression and mediate complete killing. For the heat shock promoter, its response to heat shock may be more effective during larval development than embryogenesis. Second, heat-shock induced expression of SUP-35 may not provide enough dosage to kill *C. briggsae* larvae. Further optimization of heat shock condition or choice of a different heat-shock promoter is necessary to develop an effective negative selection marker against extrachromosomal array in *C. briggsae*.

## Acknowledgments

We thank Ms Cindy Tan for logistic support and members of Zhao’s lab for helpful comments. Some strains were provided by the CGC, which is funded by NIH Office of Research Infrastructure Programs (P40 OD010440).

## Supporting information

### Figure legends for supplemental figures

**Fig. S1** Overview of plasmid components.

**Fig. S2** Identification of equivalent of *hsp-16.41* promoter in *C. briggsae*.

A. Schematics of *Steps* in identifying *C. briggsae* heat shock promoter. The *hsp-16.41* promoter sequence was derived from pMA122, which is shared by two adjacent gene pair, *hsp-16.41* and *hsp-16.2*. Genomic sequences spanning the two genes were aligned against *C. briggsae* genome “cb4” to identify their homology pairs. The cbr-HSP pair1 (grey) shares the highest similarity in coding sequence to the *C. elegans* pair, in which CBG19185 is the best hit of *hsp-16.41*. However, cbr-HSP pair2 (brown) shows the highest similarity in promoter sequence to the *C. elegans* promoter (data not shown).

B. Sequence of *C. briggsae* heat shock promoter tested in this study.

